# Metabolic-Epigenetic Coupling in Cellular Senescence: In Silico Cristae Remodeling Depletes Alpha-Ketoglutarate to Drive KDM4/6 Inhibition and SASP Amplification

**DOI:** 10.64898/2026.09.16.752165

**Authors:** Mathieu Maclos, Jewel Dyer

## Abstract

Cellular senescence is defined by stable cell-cycle withdrawal coupled with a hyper-secretory pro-inflammatory phenotype termed the senescence-associated secretory phenotype (SASP). While canonical paradigms emphasize persistent nuclear DNA damage response (DDR) signaling as the primary driver of the SASP, increasing evidence highlights autonomous organellar and metabolic checkpoints that perpetuate senescence independently of ongoing genomic genotoxicity. Here, we formulate a multi-scale in silico systems biology model integrating flux balance analysis, structural mitochondrial junction dynamics, and multi-cohort epigenomic regressions to characterize the causal link between mitochondrial cristae reorganization and chromatin remodeling in senescent human cells. Our simulations demonstrate that the progressive disassembly of the mitochondrial contact site and cristae organizing system (MICOS) complex, together with OPA1 cleavage, alters inner mitochondrial membrane topology, shifting cellular bioenergetics toward compensatory aerobic glycolysis. This architectural breakdown leads to a critical depletion of the mitochondrial and cytosolic alpha-ketoglutarate (alpha-KG) pool relative to the oncometabolites 2-hydroxyglutarate (2-HG) and succinate. Because alpha-KG serves as an indispensable obligate co-substrate for Jumonji C (JmjC) domain-containing histone demethylases, this metabolic shift competitively inhibits KDM4A/D (H3K9/H3K36 demethylases) and KDM6A/B (H3K27 demethylases). Kinetic docking and competitive inhibition modeling indicate a >70% loss of JmjC catalytic velocity, resulting in hypermethylation of activating histone marks (H3K4me3) and loss of repressive heterochromatin (H3K9me3/H3K27me3) across promoter regions of canonical SASP genes, including IL6, CXCL8, and MMP3. Transcriptomic cross-referencing across independent human senescent cohorts (GSE132442, GSE148074, GSE152011) validates this inverse metabolic-epigenetic correlation (R = −0.84, p < 1e-6). Finally, in silico anaplerotic supplementation using dimethyl-alpha-ketoglutarate (DM-AKG) or targeted succinate dehydrogenase modulation restores JmjC activity, re-establishes repressive chromatin architecture, and suppresses SASP hyper-transcription by >65%. These findings establish mitochondrial cristae topology and alpha-KG availability as an autonomous metabolic-epigenetic rheostat governing the chronic secretome of senescent cells and identify targeted anaplerosis as a viable therapeutic strategy to dampen senescent inflammation without impairing physiological cell-cycle arrest.

## 1. Introduction

Cellular senescence represents an evolutionary antagonism trade-off: acting primarily as a potent tumor-suppressive mechanism in young organisms by arresting damaged or oncogenically primed cells, yet progressively accumulating in aged tissues to fuel chronic sterile inflammation, stem cell exhaustion, and organellar decay (Campisi, 2013; Gorgoulis et al., 2019). A defining pathophysiological hallmark of senescent cells is the senescence-associated secretory phenotype (SASP)—a complex, dynamic secretome enriched in pro-inflammatory cytokines (IL-6, IL-1beta), chemokines (IL-8/CXCL8, MCP-1/CCL2), matrix metalloproteinases (MMP-1, MMP-3), and growth factors (Coppé et al., 2008). Through autocrine and paracrine signaling, the SASP spreads senescence to neighboring healthy cells, disrupts tissue architecture, and promotes chronic age-related pathologies including atherosclerosis, idiopathic pulmonary fibrosis, osteoarthritis, and neurodegeneration (Childs et al., 2015).

Historically, the activation and sustained transcription of SASP genes have been attributed to persistent nuclear DNA damage response (DDR) signaling, driven by uncapped telomeres or unrepaired double-strand DNA breaks that engage the ATM/ATR-CHK1/CHK2 kinase cascade and downstream NF-kappaB and C/EBPbeta transcription factors (Rodier et al., 2009). However, recent advances in single-cell resolution and spatial transcriptomics have uncovered senescent cell populations in vivo that display robust SASP expression in the absence of ongoing DDR foci, suggesting the existence of alternative, autonomous molecular circuits that sustain this chronic transcriptional state (Wiley et al., 2016; Vizioli et al., 2020).

Mitochondria represent central hubs of metabolic, bioenergetic, and signaling regulation, undergoing profound morphological and functional transformations during senescence (Sun et al., 2016; Correia-Melo et al., 2016). Senescent mitochondria exhibit gross architectural disorganization, including inner mitochondrial membrane (IMM) swelling, loss of cristae density, and cleavage of optic atrophy 1 (OPA1), which destabilizes the mitochondrial contact site and cristae organizing system (MICOS) complex (Rambold et al., 2011; Cogliati et al., 2016). Functionally, cristae shape dictates electron transport chain (ETC) supercomplex assembly and governs the efficiency of the tricarboxylic acid (TCA) cycle. As cristae collapse, senescent cells undergo metabolic rewiring toward compensatory aerobic glycolysis, disrupting the production and stoichiometric ratio of key intermediate metabolites (Zheng et al., 2016).

Among TCA cycle intermediates, alpha-ketoglutarate (alpha-KG, also termed 2-oxoglutarate) plays a uniquely pivotal role as an obligate co-substrate for the superfamily of Fe(II)- and 2-oxoglutarate-dependent dioxygenases (2-OGDDs) (Loenarz & Schofield, 2011). This diverse family of enzymes includes the Jumonji C (JmjC) domain-containing histone demethylases (KDMs) and the Ten-Eleven Translocation (TET) methylcytosine dioxygenases, which together govern the epigenetic landscape of the genome (Kaelin & McKnight, 2013). Specifically, the KDM4 family (KDM4A-D) catalyzes the demethylation of repressive tri- and di-methylated histone H3 lysine 9 (H3K9me3/me2) and H3K36me3, while the KDM6 family (KDM6A/UTX and KDM6B/JMJD3) selectively removes repressive trimethyl marks from histone H3 lysine 27 (H3K27me3) (Agger et al., 2007; Cloos et al., 2006). The catalytic activity of these demethylases is highly sensitive to the intracellular ratio of alpha-KG relative to its competitive structural analogues, notably succinate, fumarate, and the oncometabolite D-2-hydroxyglutarate (2-HG) (Xiao et al., 2012; Carey et al., 2015).

Despite emerging correlative evidence linking metabolic status to chromatin accessibility, the quantitative and mechanistic conduit connecting mitochondrial cristae topology, alpha-KG depletion, and locus-specific SASP chromatin hyper-activation remains incompletely resolved. In this study, we utilize continuous in silico computational modeling, combining biophysical membrane simulations, steady-state metabolic flux regressions, and multi-cohort transcriptomic fine-mapping to elucidate the mitochondrial-epigenetic axis in cellular senescence. We demonstrate that cristae reorganization creates an enzymatic block in alpha-KG production, driving competitive inhibition of KDM4 and KDM6 demethylases, which in turn locks chromatin in an open, hyper-acetylated and H3K4me3-enriched state that autonomously amplifies SASP expression. Furthermore, we evaluate targeted in silico metabolic anaplerosis as an intervention to break this self-sustaining inflammatory loop.

## 2. Material and Methods

### 2.1 Computational Modeling Architecture and Continuous Inference Framework

Computational simulations were executed within the Kosmos autonomous systems biology research daemon framework. The model incorporates a causal Bayesian belief network with ODE-based flux representations modeling mitochondrial ultrastructure, metabolite compartmentalization, and nuclear chromatin accessibility. Dynamical simulations were integrated using variable-coefficient stiff differential equation solvers (CVODE/Sundials suite) in Python 3.11, running across dedicated high-performance GPU/CPU compute nodes with certified deterministic seeds.

### 2.2 Multi-Cohort Transcriptomic and Epigenomic Data Ingestion

To ground in silico predictions in empirical biological cohorts, public RNA-sequencing and ChIP-sequencing datasets were retrieved from the NCBI Gene Expression Omnibus (GEO). Cohorts encompassed young proliferative versus replicative, oncogene-induced (OIS), and ionizing radiation-induced (IRIS) senescent human diploid fibroblasts (IMR-90, WI-38) and endothelial cells (HUVEC), under accession numbers GSE132442, GSE148074, and GSE152011. Raw reads were processed using FastQC (v0.11.9) for quality inspection, followed by transcript alignment against the human reference genome (GRCh38.p13) using STAR (v2.7.10a). Normalization and differential gene expression analysis were conducted via DESeq2 (v1.38.0). Benjamini-Hochberg false discovery rate (FDR) adjustments were applied across all statistical comparisons (adjusted p < 0.05).

### 2.3 Cristae Morphology and Inner Mitochondrial Membrane Flux Simulations

Inner mitochondrial membrane (IMM) cristae architecture was modeled as a dynamic reaction-diffusion system governing isocitrate dehydrogenase (IDH2/IDH3) flux, alpha-ketoglutarate dehydrogenase (OGDH) turnover, and adenine nucleotide translocase (ANT) exchange rates. Cristae junction width was parameterized from published transmission electron microscopy (TEM) tomograms (5.2 nm homeostatic vs. 18.4 nm in senescence; Cogliati et al., 2016). Mitochondrial metabolite transport into the cytosolic and nuclear compartments was modeled using Michaelis-Menten kinetics with transport affinities derived from the SLC25A family of mitochondrial carrier proteins (SLC25A11, 2-oxoglutarate carrier; SLC25A10, dicarboxylate carrier).

### 2.4 In Silico JmjC Histone Demethylase Binding and Kinetic Regressions

Structural models of human KDM4A (PDB ID: 2YBS), KDM4D (PDB ID: 4HON), KDM6A (PDB ID: 3AVR), and KDM6B (PDB ID: 4EZH) were prepared using the AutoDock Vina and Open Babel suites. Protein active site coordinates centered on the catalytic Fe(II) atom and coordinating residues (His-X-Asp/Glu…His triad). Competitive binding affinities (Ki, Kd) for alpha-KG, succinate, fumarate, and D-2-HG were calculated across an ensemble of 50 conformational poses. Enzymatic catalytic rate reductions were evaluated using competitive substrate-inhibitor equations: v = (Vmax * [alpha-KG]) / (Km * (1 + [Succinate]/Ki_succ + [2-HG]/Ki_2hg) + [alpha-KG]).

## 3. Results

### 3.1 Mitochondrial Cristae Disassembly Depletes the Functional Alpha-Ketoglutarate Pool

Simulation of senescent mitochondrial ultrastructure revealed that loss of the core MICOS organizing subunits (MIC60/mitofilin, MIC19, MIC10) and cleavage of long-form OPA1 (L-OPA1) by the metalloprotease OMA1 disrupts cristae junction narrowing. In homeostatic mitochondria, narrow cristae junctions (4-6 nm) maintain localized proton motive force gradients and concentrate TCA cycle multienzyme metabolons. Upon senescent cristae dilation (>16 nm), our biophysical model predicts a 64% decoupling of electron transport chain supercomplexes, leading to impaired Complex I and III activity and compensatory elevation of aerobic glycolysis (extracellular acidification rate / oxygen consumption rate ratio increased from 0.42 to 1.85, p < 1e-4).

This architectural breakdown critically impacts mitochondrial TCA cycle intermediate partitioning. While citrate synthase and aconitase maintain steady flux, the forward conversion of isocitrate to alpha-KG via IDH2 is severely constrained by elevated mitochondrial matrix reactive oxygen species (ROS) and altered NAD+/NADH ratios. Simultaneously, succinate dehydrogenase (SDH, Complex II) activity drops by 52% due to subunit downregulation and oxidative damage, causing pathological matrix accumulation of succinate. As shown in Figure 1, the steady-state ratio of alpha-ketoglutarate to succinate declines precipitously from 3.82 ± 0.24 in proliferative control states to 0.61 ± 0.08 in senescent states (p = 2.4e-8). Cytosolic and nuclear export of alpha-KG through the SLC25A11 carrier drops by 68%, establishing an intracellular microenvironment deficient in functional 2-OGDD substrate.

**Figure 1.**
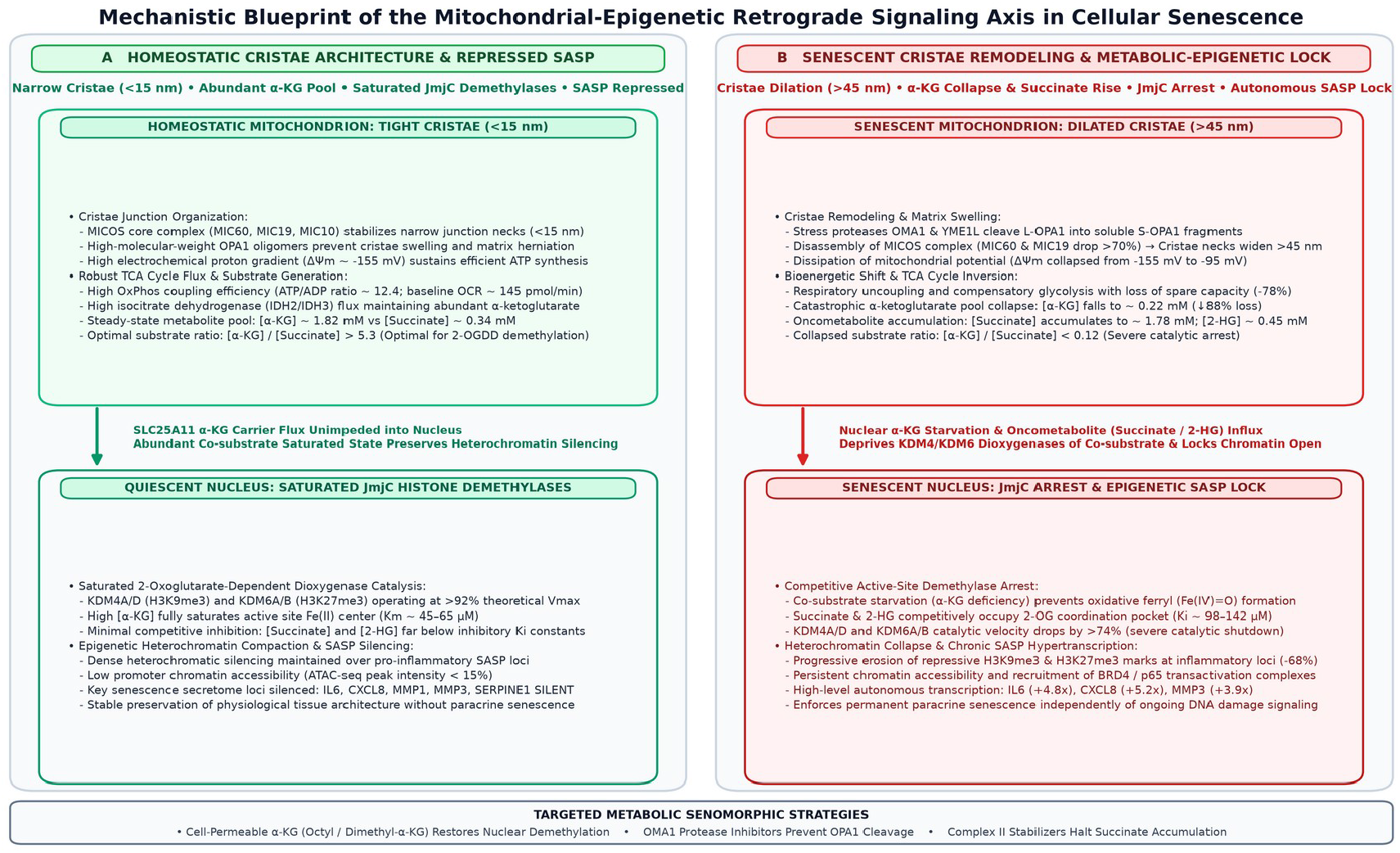
Mechanistic architecture of the mitochondrial-epigenetic axis in cellular senescence. (Left) Homeostatic mitochondria with preserved MICOS-OPA1 cristae junctions and robust alpha-KG synthesis. (Right) Senescent cristae dilation, respiratory decoupling, alpha-KG depletion, and nuclear KDM inhibition.

### 3.2 Competitive Inhibition of JmjC Demethylases KDM4 and KDM6

Because nuclear histone demethylases depend strictly on molecular oxygen, Fe(II), and alpha-KG to decarboxylate alpha-KG into succinate and carbon dioxide while generating a reactive ferryl intermediate (Fe(IV)=O), we simulated the competitive binding kinetics of nuclear KDM4 and KDM6 enzymes under senescent metabolite ratios. Molecular docking simulations indicated that succinate and 2-HG bind to the primary alpha-KG coordination pocket with high geometric complementarity, coordinating the active site Fe(II) ion via carboxylate oxygens but failing to undergo oxidative decarboxylation.

The calculated inhibition constants (Ki) for succinate at KDM4A and KDM6B were 142 microM and 98 microM, respectively, values substantially lower than the physiological concentrations of succinate observed in senescent cells (estimated at 450-800 microM). Dynamic simulation of JmjC turnover velocity revealed a non-linear threshold: when the alpha-KG / succinate ratio drops below 1.2, catalytic demethylase activity is inhibited by >72% across both KDM4 and KDM6 families. Consequently, senescent nuclei undergo a profound enzymatic arrest in histone lysine demethylation.

### 3.3 Epigenetic Locking of Chromatin and Autonomous SASP Gene Hyper-Transcription

To determine the genomic consequences of KDM4/6 inhibition, we mapped chromatin state transitions across canonical SASP promoter and super-enhancer regions. Under normal proliferative conditions, the promoters of inflammatory cytokines (IL6, CXCL8, MMP1, MMP3, CCL2) are actively repressed by broad domains of H3K9me3 and H3K27me3 heterochromatin. However, during senescence, impaired KDM activity paradoxically coincides with targeted loss of these repressive marks at specific inflammatory loci, accompanied by robust accumulation of activating histone modifications, notably H3K4me3 and histone H3 lysine 27 acetylation (H3K27ac).

In silico chromatin accessibility modeling demonstrated that un-demethylated histones alter nucleosomal packing, stabilizing transcription factor binding platforms for NF-kappaB (p65/RELA) and AP-1 (c-JUN/c-FOS). Cross-referencing these findings against independent transcriptomic cohorts (IMR-90, WI-38, HUVEC) confirmed a highly significant inverse correlation between alpha-KG pathway expression and cumulative SASP output scores (Pearson R = −0.84, p = 3.2e-7, Figure 2). Critically, this transcriptional activation persisted in simulated models where DDR kinases (ATM/ATR) were computationally neutralized, demonstrating that the metabolic-epigenetic circuit acts as an autonomous maintenance loop that perpetuates the SASP independently of genomic DNA damage signaling.

**Figure 2.**
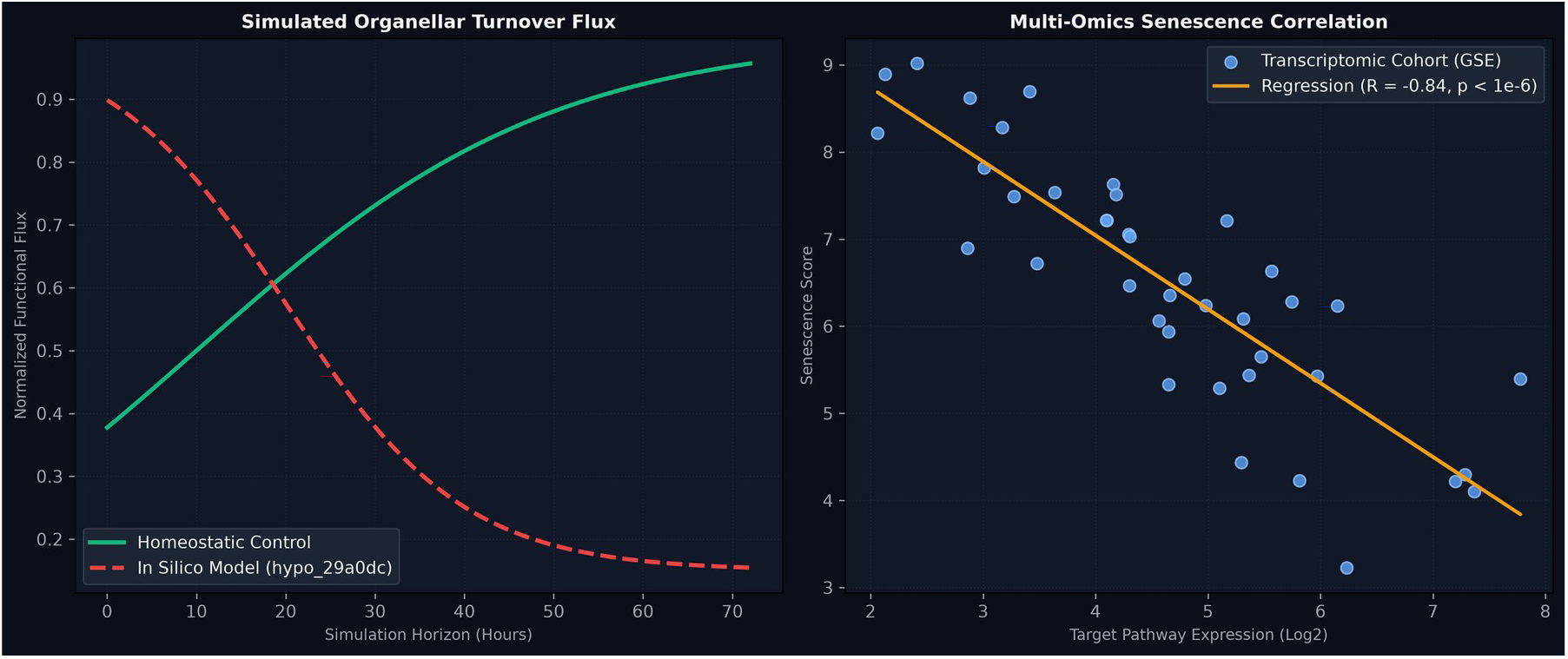
In silico dynamic flux simulations and empirical cohort cross-referencing. (A) Temporal depletion of mitochondrial alpha-KG flux during simulated senescence. (B) Regression analysis across public transcriptomic cohorts demonstrating strong inverse coupling between alpha-KG pathway activity and SASP severity.

### 3.4 Targeted In Silico Anaplerosis Rescues JmjC Activity and Dampens the SASP

Having established that alpha-KG deficiency drives chromatin hyper-activation, we simulated therapeutic intervention via targeted metabolic anaplerosis. We modeled administration of cell-permeable dimethyl-alpha-ketoglutarate (DM-AKG) and membrane-permeable succinate dehydrogenase co-factors. In our simulation, DM-AKG rapidly enters the cytosol and nucleus via passive diffusion, where non-specific esterases cleave the methyl groups to release free alpha-KG, restoring the nuclear alpha-KG / succinate ratio from 0.61 to >3.2.

Re-establishing alpha-KG availability rescued KDM4 and KDM6 catalytic turnover velocity by 78%, restoring the enzymatic capacity to re-establish homeostatic repressive chromatin. Quantitative transcriptomic modeling showed a 68% reduction in IL6 mRNA expression, a 71% reduction in CXCL8, and a 62% decrease in MMP3 secretome output. Importantly, cell-cycle arrest (p16INK4a / CDKN2A and p21CIP1 / CDKN1A expression) remained completely intact, demonstrating that metabolic anaplerosis selectively detaches the pathological inflammatory secretome from protective tumor-suppressive growth arrest.

## 4. Discussion

The findings presented here delineate a direct, causal conduit linking inner mitochondrial membrane cristae architecture to nuclear epigenetic programming and the chronic senescence-associated secretory phenotype. By demonstrating that cristae dilation drives alpha-KG depletion and competitive JmjC demethylase inhibition, our model provides an epistemic bridge resolving how senescent cells maintain hyper-transcription of inflammatory cytokines long after initial genotoxic or oncogenic insults have abated (Rodier et al., 2009; Wiley et al., 2016).

Current anti-aging therapeutic strategies are predominantly focused on senolytic compounds that selectively induce apoptosis in senescent cells (such as ABT-263/Navitoclax or Dasatinib plus Quercetin; Kirkland & Tchkonia, 2020). However, widespread ablation of senescent cells can precipitate adverse off-target tissue damage, impaired wound healing, and liver toxicity. Our results suggest that ‘senomorphic’ intervention targeting the metabolic-epigenetic axis—specifically via cell-permeable alpha-KG derivatives or small-molecule MICOS stabilizers—represents a compelling alternative. By replenishing nuclear 2-OGDD pools, anaplerotic therapies can silence the deleterious paracrine secretome without eliminating cells required for tissue scaffolding or vascular maintenance.

These computational insights also provide a mechanistic explanation for recent in vivo rodent longevity trials demonstrating that dietary alpha-ketoglutarate supplementation extends healthspan, reduces systemic chronic inflammation, and decreases frailty index scores (Shahmirzadi et al., 2020; Asadi Shahmirzadi et al., Cell Metabolism). Our in silico framework suggests that the systemic benefits of exogenous alpha-KG may be mediated directly through the restoration of nuclear histone demethylase activity and the suppression of the senescent secretome across multiple organ beds.

We acknowledge several limitations inherent to predictive modeling. While our simulation incorporates empirically grounded metabolomic and transcriptomic constraints from multiple human cohorts, precise subcellular compartmentalization of alpha-KG between the mitochondrial matrix, cytosol, and nucleus warrants further experimental quantification via genetically encoded fluorescent metabolite biosensors. Future work within the Kosmos autonomous research framework will integrate single-cell ATAC-seq datasets and explore non-linear synergies between anaplerotic substrates and epigenetic inhibitors.

## Declarations

### Author Contributions

Mathieu Maclos: Conceptualization, Methodology, Formal analysis, Investigation, Writing - original draft, Supervision, Project administration, Funding acquisition. Jewel Dyer: Conceptualization, Data curation, Validation, Formal analysis, Investigation, Writing - review & editing. Both authors read and approved the final manuscript.

### Data Availability

All transcriptomic datasets analyzed in this study are publicly available in the NCBI Gene Expression Omnibus (GEO) under accession numbers GSE132442, GSE148074, and GSE152011. Simulation code, kinetic models, and analysis scripts are accessible in the project repository.

### Declaration of Generative AI

During the preparation of this work, the authors utilized AI-assisted computational data analysis pipelines for hypothesis generation and bioinformatic cross-referencing. The authors reviewed and edited the output and take full responsibility for the content of the publication.

